# Relaxed purifying selection is associated with an accumulation of transposable elements in flies

**DOI:** 10.1101/2024.01.23.576885

**Authors:** Vincent Mérel, Théo Tricou, Nelly Burlet, Annabelle Haudry

## Abstract

Although the mechanisms driving the evolution of genome size are not yet fully understood, one potentially important factor is the dynamics of the accumulation of mobile selfish genetic elements called transposable elements (TEs). Since most of these sequences are neutral or slightly deleterious, a negative correlation between genome size and selection efficacy is expected. However, previous studies based on empirical data from closely related species with contrasting life history traits (thought to result in contrasting levels of selection efficacy) have yielded inconsistent results, leaving this issue controversial. In this study, we perform the first large-scale analysis of the effect of drift on genome size evolution, without any prior assumption about the amount of drift in each sampled species. We reconstructed a phylogeny based on whole-genome data (2,242 genes) for 77 Drosophilid species to examine correlations between genome size, TE content, and measures of selection efficacy (especially using *dN/dS* ratios of non-synonymous to synonymous divergence). We highlight a strong phylogenetic inertia in genome size and confirm that TEs are the major components of genome size. Using an integrative approach that controls for shared history, we find that genome-wide *dN/dS* are strongly positively correlated with genome size and TE content, particularly in GC-poor genes. This study highlights the critical importance of controlling for heterogeneity in base composition when testing the controversial correlation between evolutionary rates and genome size. Furthermore, our review of previous studies reveals that the absence of evidence for TE accumulation in association with increased genetic drift may be attributed to a secondary effect of changes in life history traits on TE dynamics. In conclusion, this work provides evidence for TE proliferation in fly genomes when purifying selection is reduced and genetic drift increases, shedding new light on the role of transposable elements and genetic drift in the evolution of genome architecture.

## 1 Introduction

Eukaryotic species exhibit a surprisingly wide range in genome size: the genome of the parasitic nematode *Pratylenchus coffeae* is approximately 0.019 Gbp (Gregory, 2005), whereas that of the fork fern *Tmesipteris oblanceolata* was recently estimated to reach 160 Gbp (Fernández et al., 2024). The accumulation of repeated elements is thought to be the main driver of genome size evolution in Eukaryotes. Especially, genome size has been found strongly positively correlated to the amount of Transposable Elements (TEs) in various taxa (Biémont and Vieira, 2006; Tenaillon et al., 2010; Elliott and Gregory, 2015). TEs are selfish genetic elements (Orgel and Crick, 1980), mostly neutral or deleterious to their host, that persist and even proliferate, by copying and pasting themselves to other genomic locations. Genome size therefore reflects the dynamics of TE accumulation, depending on rate of TE insertion and deletion. As an example, the expansion identified in the salamander genome results from an increased proliferation of LTR retrotransposons (Sun et al., 2012).

Due to the deleterious effects of TEs, purifying selection is expected to prevent TE fixation and thus to maintain small genomes. Effective population size (*Ne*) which affects selection efficacy (*Nes*, with *s* the selection coefficient), has been proposed as a key parameter controlling TE accumulation (Lynch and Conery, 2003). Since selection efficacy is positively correlated with *Ne*, TEs are expected to accumulate in lineages where *Ne* is reduced. In line with the *Mutational Hazard* (MH) hypothesis, Lynch and Conery (2003) found a negative correlation between synonymous diversity (an estimate of 4*Neµ, µ* being the mutation rate) and genome size across species. However, Lynch and Conery (2003) findings appear to be largely explained by shared common evolutionary history (Whitney et al., 2010), and rely on the comparison of very distant species with drastically different biology, which could act as a confounding factor (Charlesworth and Barton, 2004).

The distribution of quantitative traits among species is not independent of the length of the shared history between the considered species: closely related species tend to resemble each other more than expected by chance, due to *phylogenetic inertia* (Felsenstein, 1985). As a result, it is necessary to account for the phylogenetic relatedness of species and the resulting dependencies in comparative analyses. Approaches that explicitly account for the phylogenetic signal to correct for correlations between traits rely on stochastic models of trait evolution — typically assuming that the traits jointly evolve according to a multivariate Brownian process running along the lineages of the phylogeny. The covariance matrix of this multivariate process is then estimated either by maximum likelihood (e.g. phylogenetic generalised least squares PGLS, (Martins and Hansen, 1997)) or by Bayesian inference (Pagel and Harvey, 1988), accounting for phylogenetic dependencies in the estimation of the correlations between traits. Methods at the intersection between the classical comparative method and molecular phylogenetics (Lartillot and Poujol, 2011; Lartillot, 2014) have suggested that a substantial amount of information about ancestral traits could be obtained from the evolution of genetic sequences. The fundamental idea is as follows: (1) detailed patterns of genetic sequence evolution can be fairly accurately inferred along a phylogeny; (2) the variation in evolutionary rate along the branches of the tree can be inferred; (3) more fundamentally, these evolutionary rates and parameters can be seen as continuous traits and modelled as Brownian processes. All this can be achieved in a single Bayesian probabilistic model, combining the molecular phylogenetic tree and the comparative method dimensions of the question.

Beyond the genetic diversity used in Lynch and Conery (2003), a variety of traits could be related to *Ne*. According to the nearly neutral theory of evolution, smaller *Ne* should lead to an accumulation of slightly deleterious mutations, and thus to a higher ratio of non-synonymous over synonymous divergence, or *dN/dS*, which is used as a measure of selection efficacy on protein sequence (Castillo et al., 2011; Bolívar et al., 2019). Several life history traits have previously been reported to affect *Ne* (eg. maturity or longevity in mammals (Lartillot and Delsuc, 2012) or mating system in plants (Pollak, 1987)). It is therefore possible to test the effect of a reduction in *Ne* on the genome architecture using comparisons among closely related species with contrasted life history traits. Besides, population genetic theory predicts that selection on codon usage should be much weaker than selection acting on the protein sequence (typically on the order of *4Nes* ≈ 1, *s* being the selection coefficient in favour of optimal codons). Since weakly deleterious mutations are most susceptible to fixation when *Ne* is reduced (McVean and Charlesworth, 1999), codon usage bias metrics may provide alternative estimates of selection efficiency. While some fruit flies with large *Ne* show higher codon usage bias than their relatives (Haddrill et al., 2010)),Kessler and Dean (2014) found no direct relation between codon usage bias and *Ne* variation in mammals. Indeed, codon usage bias results from a complex balance between mutation patterns, selection, recombination (via GC-biased gene conversion) and genetic drift Parvathy et al. (2022). It remains an open question whether codon usage metrics can reflect *Ne* variation in large populations such as fruit flies. To address it, we will test for potential correlation between *dN/dS* and codon usage bias measures, and developmental time in flies.

Appealing because it is based on universal principles of population genetics, the MH hypothesis fuelled the need for empirical testing. To date, comparative studies of closely related taxa have reported conflicting results. Some have shown an accumulation of TEs resulting from a reduction of *Ne*, as predicted by the MH theory: in ectomycorrhizal fungi in symbiotic fungi (Hess et al., 2014); in subterranean *Aselloidea* (Lefébure et al., 2017); in polyploid plants (Baduel et al., 2019); in eusocial snapping shrimps and termites (Chak et al., 2021; Korb et al., 2015); in invasive populations of *D. suzukii* (Mérel et al., 2021)—, others found no significant excess in TE abundance in species with reduced *Ne* compared to their close relatives —in asexual daphnia (Schaack et al., 2010; Jiang et al., 2017), primroses (Ågren et al., 2015), arthropods (Bast et al., 2016) or yeast populations (Bast et al., 2019); in selfing *Caenorhabditis* nematodes (Fierst et al., 2015); in eusocial bees (Kapheim et al., 2015), or hymenoptera (Ardila-Garcia et al., 2010).

*Drosophila* flies show a three-fold variation in genome size, reflecting variation in TE content (Sessegolo et al., 2016). Differences in life history traits and demography were reported (Markow and O’Grady, 2005), which could potentially lead to *Ne* differences. Contrary to MH’s expectations, a study of 12 distant species showed that greater levels of purifying selection were rather associated with greater euchromatic TE abundance, using *dN/dS* as a proxy of *Ne* (Castillo et al., 2011), letting the genome size variation puzzling. Analyses by Sessegolo et al. (2016) revealed that strong phylogenetic inertia partially explains genome size differences, with a sampling bias towards small genome species closely related to *D. melanogaster*, which could be easily corrected with a more balanced sample across the Drosophilid phylogeny. To date, the main drivers of repeatome expansion in fly genomes remain to be identified, especially in regard to the respective roles of drift and selection.

Here, we investigate genome size, TE content and selection efficacy in 82 flies lineages from 77 species to gain insight into genome size evolution. Our study includes historical *dN/dS* reconstruction along with ancestral genome size and TE content based on interspecific divergence and evolutionary rate using a Bayesian method accounting for phylogenetic relationships, directly assessing intrinsic correlations between those variables.

## 2 Results

### 2.1 Thirty eight novel fly assemblies

To study the evolution of genome size, TE content and effective population size in flies we focused on Next Generation Sequencing (NGS) data for 82 Drosophilid lineages. This dataset includes 77 species from *Drosophila* and *Zaprionus* genus, with a particular focus on Drosophila and Sophophora subgenus. It spans 25–40 million years of divergence (Obbard et al., 2012) (supplementary table S1). Forty four already available and 40 new samples were selected in order to obtain a large set of species representing the diversity of the clade (supplementary table S2,3), resulting in a balanced sample of species along the phylogeny (at least three species were sampled per Drosophila group or Sophophora subgroup based on taxonomy). For each of the 40 new samples we constructed genomic assemblies using short-insert paired-end reads. Overall, the mean N50 of newly built assemblies is 20Kb, and range from 3 Kb to 92 Kb (supplementary table S2). We found a mean of 2,941 complete genes out of the 3,285 dipteran BUSCO genes, *i*.*e*. 89%(supplementary figure 1). Two assemblies with a high percentage of duplicated BUSCO genes (≥ 20) were not further considered (*D. algonquin* and *D. tristis*). Homemade assemblies include ten species that, to our knowledge, have not been sequenced so far *D. lucipennis, D. lutescens, D. madeirensis, D. microlabis, D. mimetica, D. pallidosa, D. paralutea, D. prostipennis, D. tsukubaensis*, according to NCBI. And four species that have been sequenced but for which no assembly was available (*D. affinis, D. helvetica, D. imaii, D. mercatorum*). These genome assemblies were used to collect orthologous genes in order to built the phylogeny and estimate genomic traits such as the *dN/dS* ratio. Although the assemblies could be quite fragmented, the number of predicted genes (using AUGUSTUS) was generally close to the annotated 13,986 coding genes of *D. melanogaster* (according to FlyBase release 112.10). This suggest that most coding regions were assembled. Furthermore, it is important to note that the estimates of genome size or TE content are independent of the quality of the assemblies (see the Methods section for details).

**Figure 1:**
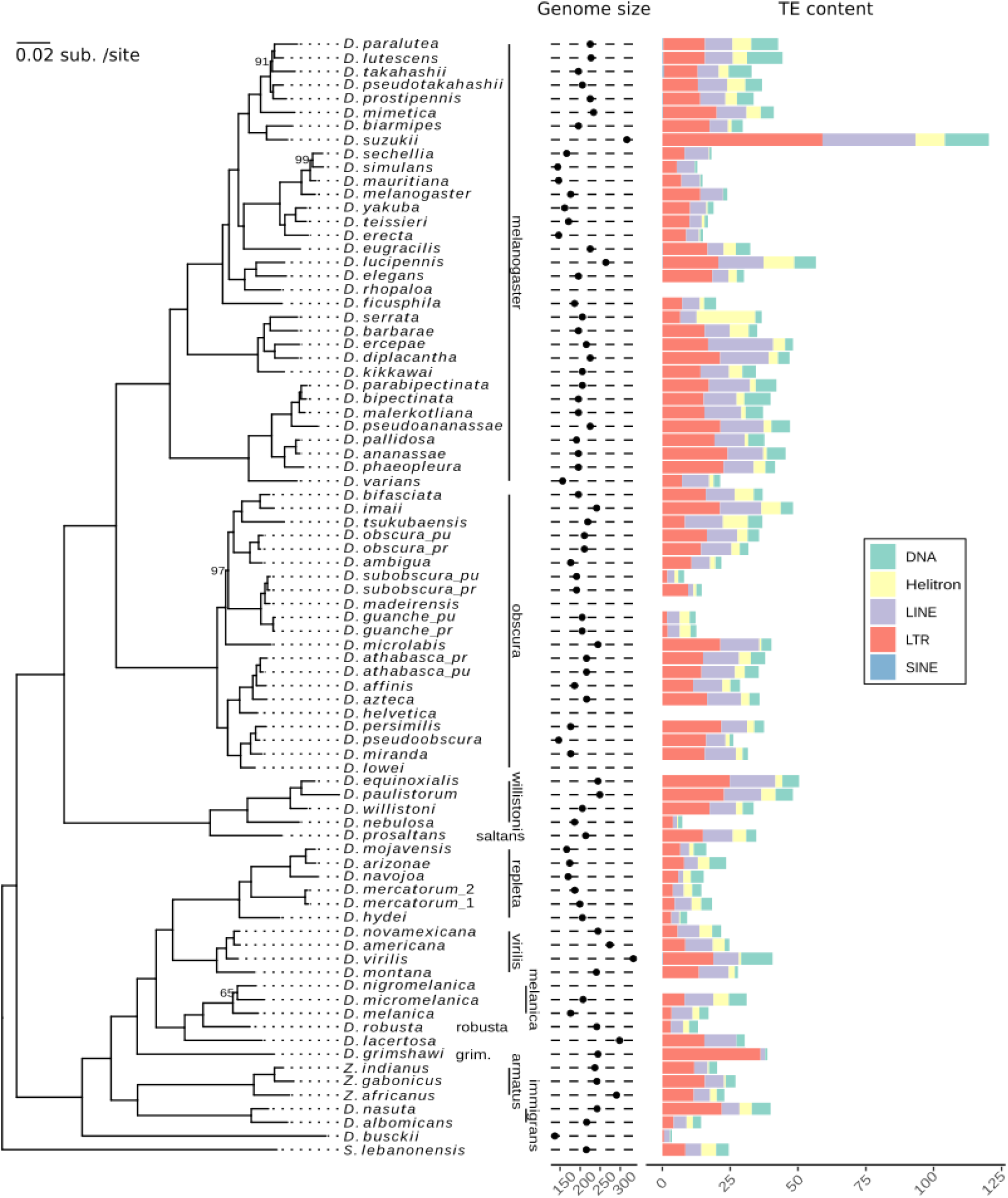
Phylogenetic tree figuring genome size and TE abundances. The tree was reconstructed using 2,242 orthologous genes and a maximum likelihood approach. Bootstraps values are indicated at node when different from 100. Taxonomic groups according to Schoch et al. (2020) are indicated on the right. Genome size (Mb) was estimated using flow cytometry, and genomic repeat content (Mb) was estimated from short reads using dnaPipeTE (Goubert et al., 2015).

### 2.2 *Drosophila* phylogeny reconstruction

We used a *de novo* clustering approach in order to identify single-copy orthologous genes present in at least 50% of the genomes (*i*.*e*. in at least 41 genomes). The resulting 2,242 nuclear orthologous genes were aligned, then concatenated to reconstruct a phylogenetic tree, using *Scaptodrosophila lebanonensis* as an outgroup (figure 1). This Maximum Likelihood (ML) phylogeny reports highly confident node bootstrap support (only four nodes with UFBoot <100) and identifies nine main clades, consistent with taxonomic Drosophila and Sophophora groups.

This tree was compared to an ASTRAL supertree, i.e. an estimation of the true species tree from the 2,242 unrooted gene trees. Our supertree ASTRAL topology is discordant with the ML topology in only four relationships, three of them being associated with the least supported nodes in the ML tree. First, *D. micromelanica* is found at the root of the *melanica* group in the ASTRAL tree, while it is *D. melanica* in the ML tree (UFBoot = 65, figure 1). Second, *D. hydei* is sister to the species group of *D. navojoa, D. arizonae, D. mojavensis* in the ASTRAL tree, while it is also sister to *D. mercatorum* lines in the ML tree (UFBoot = 100). Third, *D. microlabis* is sister to the obscura subgroup only in the ASTRAL tree, but to both the *obscura* and the *subobscura* subgroups in the ML tree (UFBoot = 97). Last, *D. takahashii* is at the root of the whole *takahashii* subgroup in the ASTRAL tree, but sister to *D. lutescens* and *D. paralutea* in the ML tree (UFBoot = 91). We compared the present topology with the Suvorov’ ML phylogenomic tree using a subsample of the 55 species shared between the two datasets (supplementary figure 2). Only two incongruencies are found, resulting in a Robinson Foulds distance metric of 8 between trees (Robinson and Foulds, 1981): (1) the sister species of *D. simulans* is *D. sechellia* in our topology while *D. simulans* groups with *D. mauritiana* in the Suvorov tree. (2) *D. ercepeae* groups in the *montium* subgroup (*melanogaster* group) while it branches at the basis of the *ananassae* subgroup (*melanogaster* group) in the Suvorov tree, as well as in Van der Linde’s tree.

**Figure 2:**
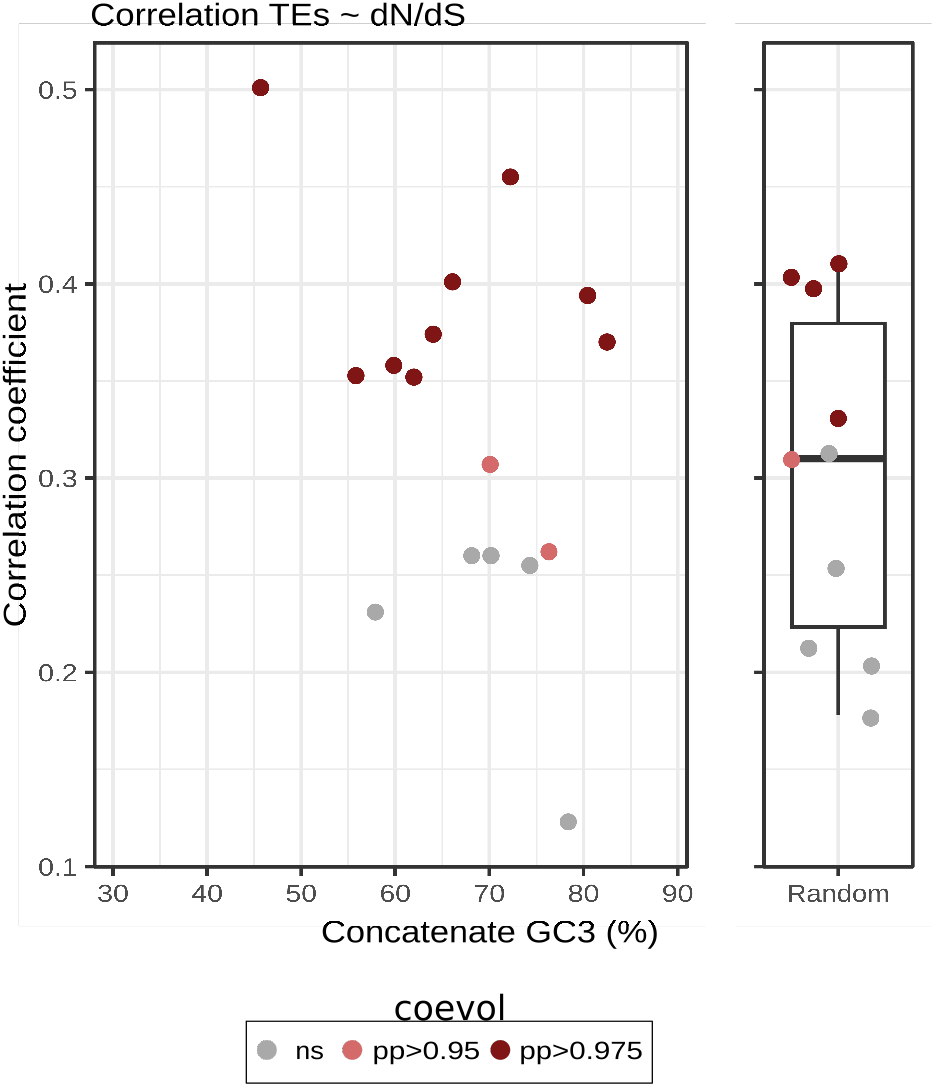
Correlation coefficients between TE abundances and *dN/dS* as a function of GC3. The dots indicate the median GC3 for each concatenate. Correlation coefficients were estimated using coevol. Strongly and very strongly supported positive correlations are indicated in light and dark red respectively (pp>0.975 and pp>0.95).

### 2.3 Strong phylogenetic inertia on genome size and TE content

We retrieved flow cytometry estimates of genome size for 77 out of our 82 samples. 54 of these estimates come from the literature (Gregory, 2005; Hjelmen et al., 2019), the remaining 23 estimations were performed for this study. The genomic size ranges from 136.92 Mb for *D. busckii* to 332.52 Mb for *D. virilis*, with a mean of 208.7 Mb (supplementary table S4, figure 1). We found a Pagel’s lambda very close to one for this trait, indicating a strong effect of shared ancestry (Pagel’s λ = 0.92, *p* = 1.64.10^−5^).

We then estimated repeat content in our **77** samples using dnaPipeTE (figure 1, supplementary table S5)(Goubert et al., 2015). Briefly, dnaPipeTE reconstructs TE sequences from the assembly of a subsample of reads (such as coverage < 1X), annotate these sequences by homology to RepBase database (Kapitonov and Jurka, 2008), and then estimates the proportion of reads mapping on each category of repeat. This proportion can be converted into a number of base pairs, based on the genome size estimate. We found the total amount of TE to range from 3.48 Mb for *D. busckii* to 120.42 Mb for *D. suzukii*. The category with the highest average abundance is LTR elements (supplementary table S6).

We compared our estimations of TE abundances with the ones obtained by Kim et al. (2021). In the mentioned study the authors estimated repeat content in 101 Drosophilid species long read assemblies using homology to sequences in online databases. Comparing abundance estimates for the 36 species common to the two studies, we found significant correlations for all TE types (*R*^2^ ranging from 0.40 to 0.46, *p*_*adj*_ < 0.05, supplementary figure 3). Note that for this comparison 1) we considered retrotransposons (estimated in Kim et al. (2021)) to correspond to the sum of LINEs and LTR elements, 2) RC to correspond to Helitrons (Kapitonov and Jurka, 2008), 3) for SINE, we were not able to test any correlation as this category was not present in Kim et al. (2021). Our results suggest a very strong phylogenetic inertia on TE content. Pagel’s λ indicate phylogenetic signal for TEs as a whole (Pagel’s λ = 0.99, *p*_*adj*_ = 1.64.10^−5^), but also for DNA, Helitron, LINEs and LTR elements alone. Only for SINEs results were not significant.

### 2.4 Genome size correlates with abundances of various repeat types

We tested pairwise correlations between genome size and abundances of the different TEs using two different methods, both controlling for phylogenetic inertia (table 1). The first one, implemented ancov (Lartillot, 2014), performs Bayesian reconstruction of traits to estimate the correlation among all traits in one single test. The second one corresponds to multiple tests of Phylogenetic Generalised Least Squares (PGLS), adjusted for multiple testing. Eleven traits were found to be significantly correlated with both methods, three private to ancov, and two to PGLS. Overall, we found genome size to be positively correlated with the total amount of TEs (ancov correlation coefficients of 0.62). However, the contribution to genome size variation was different between TE categories, with LINE, DNA and LTR as main contributors. We also found significant correlation between the abundances of various categories of transposable elements. The strongest, between DNA and LINEs, had a correlation coefficient of 0.45.

**Table 1:** Pairwise correlations between genome size and TE abundances. Correlation coefficients were estimated using ancov. Cells in light gray correspond to significant correlations (posterior probability (pp) < 0.025 or pp > 0.975). Significant PGLS are presented in bold (Benjamini-Hochberg *p*- adjusted < 0.05)

|  | GS | DNA | Helitron | LINE | LTR | SINE | TEs |
| --- | --- | --- | --- | --- | --- | --- | --- |
| GS |  | **0.55** | **0.19** | **0.56** | **0.38** | 0.03 | **0.61** |
| DNA | - |  | 0.21 | **0.45** | **0.31** | 0.28 | **0.58** |
| Helitron | - | - |  | **0.31** | -0.10 | -0.14 | **0.21** |
| LINE | - | - | - |  | **0.22** | 0.10 | **0.61** |
| LTR | - | - | - | - |  | 0.14 | **0.83** |
| SINE | - | - | - | - | - |  | 0.17 |

### 2.5 Genome size and TE content correlate with *dN/dS*

We further assessed if variations of genome size and TE content could be explained by changes in population effective size *Ne*. We used *dN/dS* as a proxy of *Ne*, and also tested some codon usage bias metrics (SCUO and ENC’) and a life history trait. Given that a larger *Ne* should lead to the fixation of optimal codons, the SCUO is expected to be positively correlated with *Ne* (with a SCUO close to 0 for low codon bias and close to 1 for strong bias). On the contrary, ENC’ should be negatively correlated with selection efficacy (with a ENC’ close to 20 for high codon bias and close to 60 for no bias). If a weaker selection explain genome size and abundance of repeats one may thus expect a negative correlation of these variables with SCUO, and a positive correlation with ENC’. We also tested whether the developmental time could reflect *Ne* in flies, as several life history traits have previously been shown to affect *Ne* (Pollak, 1987; Lefébure et al., 2017). However, this trait was only reported for 52 over the 82 lines and showed a few level of inter-specific variation (variance=8.95), with none within taxonomic groups. To our knowledge no correlation between developmental time and selection efficacy have been established in *Drosophila* so far.

We used coevol to jointly assess correlations between genome size, TE content and developmental time, and estimates of selection efficacy (Lartillot and Poujol, 2011). Given a tree and an alignment, coevol models the evolution of mentioned variables assuming brownian motion parametrized by a covariance matrix. All parameters are estimated in Bayesian framework using Monte Carlo Markov Chains. The posterior probability associated to each correlation can be interpreted as a statistical support. A posterior probability close to 1 corresponds to a strong statistical support for a positive correlation. Conversely, a posterior probability close to 0 corresponds to as strong statistical support for a negative correlation. In order to test for a different evolution for genes of contrasted GC3 (GC content at the third-codon position), *dN/dS* were estimated on alignments of three concatenates of 50 genes each: one composed of randomly sampled genes, one composed of genes with the lowest GC3, and one composed of genes with the highest GC3 (see supplementary figure 4 for GC3 distribution).

Our analysis reveals no correlation between developmental time and genome size/TE abundances (table 2). Two correlations were found using codon usage bias metrics, both for the concatenate of low GC3. For this concatenate the abundance of Helitron was found to be positively correlated with ENC’ and negatively with SCUO, suggesting a negative impact of *Ne* on Helitron proliferation. The use of *dN/dS* suggests a more predominant role of population effective size, with yet different results according to the GC3. For genes of high and random GC3, we found two and five significant correlations respectively, while all except one correlation were highly significant for low GC3 genes. The abundance of TEs as a whole was always found to be correlated with *dN/dS* with correlation coefficients ranging from 0.37 to 0.50. Only the abundance of SINE was never found significantly correlated to *dN/dS*.

**Table 2:** Correlation coefficients between genome size, TE abundances and traits related to *Ne*. Correlation coefficients were estimated using coevol. Asterisks indicate strength of support of the posterior probability to be different than 0 (** - pp>0.975 or pp<0.025 - very strong support, * - pp>0.95 or pp<0.05 - strong support).

|  |  | GS | DNA | Helitron | LINE | LTR | SINE | TEs |
| --- | --- | --- | --- | --- | --- | --- | --- | --- |
| $dN/dS$ | Random genes | 0.31* | 0.04 | 0.29* | 0.40** | 0.34* | -0.02 | 0.41** |
|  | Low GC3 genes | 0.48** | 0.37** | 0.37** | 0.41** | 0.40** | -0.18 | 0.50** |
|  | High GC3 genes | 0.25 | 0.20 | 0.25 | 0.18 | 0.35* | 0.02 | 0.37** |
| ENC' | Random genes | -0.03 | -0.05 | -0.07 | -0.01 | -0.03 | -0.03 | -0.02 |
|  | Low GC3 genes | 0.01 | 0.07 | 0.32** | 0.07 | -0.06 | -0.08 | 0.01 |
|  | High GC3 genes | -0.09 | -0.14 | -0.07 | -0.05 | -0.13 | -0.04 | -0.15 |
| SCUO | Random genes | -0.06 | -0.06 | 0.06 | 0.04 | 0.00 | -0.07 | -0.01 |
|  | Low GC3 genes | -0.11 | 0.01 | -0.28** | -0.10 | 0.06 | 0.10 | -0.06 |
|  | High GC3 genes | -0.17 | -0.13 | 0.07 | -0.09 | -0.09 | -0.13 | -0.15 |
| Developmental time |  | 0.13 | 0.06 | -0.03 | 0.06 | 0.05 | -0.06 | 0.10 |

### 2.6 Effect of intragenomic variation in GC3

We further investigated how GC3 affected correlations between *dN/dS*, genome size, and repeat abundance. We thus repeated the analysis with 16 independent concatenates of 50 genes with homogeneous GC3, and ten concatenates of 50 genes with random GC3.

A significant correlation between genome size and *dN/dS* was found in five (over ten) GC3-random concatenates and in seven (over 16) GC3-homogeneous concatenates (supplementary figure 5). The lowest GC3 concatenate showed a particularly elevated correlation coefficient (R=0.48), compared to the other GC3-homogeneous concatenates (R=0.27 in average, with significant R ranging from 0.30 to 0.35). Nevertheless, we did not found any negative correlation between the concatenate GC3 and the coefficient of correlation between genome size - *dN/dS* (*t* = −1.40, *p*_*adj*_ = 0.32).

Considering the abundance of TEs as a whole, we found more significant correlations with *dN/dS* than with genome size: five for GC3-random concatenates (over ten) and eleven (over 16) for GC3-homogeneous ones (figure 2). Once again, the lowest GC3-concatenate had a particularly elevated correlation coefficient (R=0.50) compared to the other GC3-homogeneous (in average R=0.33), and the correlation between the concatenate GC3 and the coefficient of correlation between TE content and *dN/dS* was not significant (*t* = −1.51, *p*_*adj*_ = 0.32). Investigating TE abundances per type, LINEs, LTR elements and Helitrons were found to give similar results to TEs and genome size with at least eleven (out of 26) concatenates supporting a positive impact of relaxed purifying selection on abundances (supplementary figure 6). However, no concatenate did support such an impact for SINEs, and only two for LINEs. In any case, no correlation was found between correlation coefficients and the concatenate GC3 (pearson correlation test, *p*_*adj*_ > 0.05).

For validation, we compared our estimates of *dN/dS* from coevol, which is using an integrative approach assessing simultaneously correlations over the whole tree, with an approach only considering last-branch *dN/dS*. For each concatenate (GC3-homogeneous and random), we estimated *dN/dS* for each branch of our phylogeny using *MapNH*, a mapping algorithm (Guéguen and Duret, 2018; Minin and Suchard, 2008; Dutheil et al., 2012). We verified consistency between *dN/dS* estimates from the two methods (supplementary figure S7-8): they were positively and significantly correlated for each concatenate (*p*_*adj*_ < 0.05), with *R*^2^ ranging from 0.19 to 0.85 (mean=0.52).

## 3 Discussion

We performed comparative genomics on 82 Drosophilid lineages. To do so, we generated *de novo* assemblies for 38 *Drosophila* species, 14 of which did not have their genome assembled before. Based on a clustering approach of predicted genes, we identified over 2,200 single copy orthologous genes. This allowed the reconstruction of a robust phylogeny, highly congruent with previously established ones (Van Der Linde et al., 2010; Suvorov et al., 2020), including ten supplementary species. We also estimated genome size using flow cytometry for 23 lineages. These new genomes and genomic data provide to the community valuable resources in species closely related to the reference species *D. melanogaster*, and open the possibility to further investigations.

### 3.1 *Drosophila* phylogeny

The ML topology we report here is highly congruent with previous *Drosophila* tree reconstructions of Suvorov et al. (2020) and Van Der Linde et al. (2010), with only two incongruencies. First, the sister species of *D. simulans* is *D. sechellia* in our topology while *D. simulans* groups with *D. mauritiana* in the Suvorov tree. However, Suvorov and co-authors also identified *D. sechellia* as sister species of *D. simulans* on their ASTRAL topology and suggested that this is the most likely according to a focus on low-recombining regions, which are less prone to incomplete lineage sorting. Second, *D. ercepeae* groups in the *montium* subgroup (*melanogaster* group) while it branches at the basis of the *ananassae* subgroup (*melanogaster* group) in the Suvorov tree, as well as in Van der Linde’s tree. Because both Suvorov and the present trees are based on thousands genes (some most likely in common), it does not seem reasonable to suggest horizontal transfer events to explain the discrepancy. There are several possible explanations for this consequent change in the place of *D. ercepeae*. It could either be due to contamination (mislabelled fly species) or a subsample effect. Indeed, our topology was compared to the one of Suvorov et al. (2020) on the basis of shared species, representing subsamples of original datasets; however each alignment trimming parameters and model of substitution are chosen for a whole dataset, a subsample tree depends to some extent of the original one. According to this hypothesis, the discrepency in the place of *D. ercepeae* would indicate that this species may have a complex evolutionary history.

### 3.2 TE content evolution mostly explains genome size variation within flies

Overall, genome size and TE content are strongly positively correlated within flies, mostly due to LINEs, DNA and LTR elements. As LINEs and DNA elements are generally shorter than LTR elements in *Drosophila* (see Mérel et al. (2020) for a review), their predominant role in genome size evolution is likely to reflect a larger variation in copy number. However, for LINEs and LTR elements, the correlations with genome size may also be blurred by a slight underestimation of elevated content. For species where Kim et al. (2021) found retrotransposons to be particularly abundant in long reads based assemblies, our estimates were lower (for example *D. paulistorum* with 71 Mb versus 36 Mb). This is likely to reflect the difficulty of correctly assembling repeats from a subsample of short reads when they are too numerous and divergent. Importantly, it may also negatively affect the moderate relationship between genome size and TE abundance (R ∼ 0.6).

In any case, TE content appears to be the main driver of genome composition at a rather short evolutionary scale (here ∼ 60 My), as it was previously reported for Eukaryotes (Kidwell, 2002). TEs *per se* may play a fundamental role in the differentiation among genomes, as they represent a “junk pool” of large standing mutations and might potentially act on species divergence. However, we can note that we only detected an average (for a fly) TE content for *D. virilis*, which has the largest genome reported so far in *Drosophila* species. As the genome of *D. virilis* consists of more than 40% of a few related 7-bp satellites (Flynn et al., 2020), the additional DNA accumulated in its genome does not correspond to TEs and is therefore not represented here (as we specifically chose to focus on TEs).

Our results show a strong effect of shared ancestry on both genome size and TE content indicating that closely related species tend to bear more similar repeat content (and thus genome size) compared to distant species, in agreement with results in 26 fly species reported by Sessegolo et al. (2016). In addition to strong phylogenetic inertia, we found that both genome size and TE content vary within Sophophora (melanogaster, obscura, willistoni and saltans groups) and Drosophila subgenus (including repleta, virilis, melanica, armatus and immigrans groups). Genome size and TE content also vary within groups of Sophophora or Drosophila subgenus. In particular, *D. suzukii* experienced an increase in genome size associated with an accumulation of diverse repeated elements especially since its divergence from *D. biarmipes*, as it was previously reported by Sessegolo et al. (2016). Furthermore, in the obscura group, *D. guanche* (n=2) and *D. subobscura* (n=2) lineages exhibit less TEs than close relatives, such as *D. obscura* (n=2) or *D. imaii* (n=1).

The fact that TE content and genome size vary within groups of species suggests that independent changes in TEs accumulation dynamics occurred throughout the evolutionary history of flies. Such interspecific variation in genomic traits, associated with potential variation in *Ne* across the closely related species suggests that our dataset offers a great opportunity to test whether such variation in TE content (hence in genome size) could be associated with variation in selection efficacy.

### 3.3 Which genomic trait best reflects variation in genetic drift?

We used a phylogenetic approach integrating ancestral trait reconstruction to estimate the dynamic relationship between genomic traits expected to be related to *Ne* and genome size and TE content. Using randomly sampled genes, we found a significant positive correlation between the TE content and the *dN/dS* ratio, whereas no significant association was detected between genomic composition (TE content nor genome size) with the other traits related to *Ne*, namely codon usage bias statistics (ENC’ or SCUO) or a life history trait (the developmental time). These results may raise the question of which genomic traits are good proxies of *Ne* at the evolutionary scale we are looking at. While species life history traits affect diversity levels and *dN/dS* within mammals (Lartillot and Delsuc, 2012) or Metazoa (Romiguier et al., 2014), no relationship was found between developmental time and *dN/dS* at the *Drosophila* scale. One pitfall of this result relies on the absence of within group variation on this trait, making its potential effect corrected for jointly with the phylogenetic relationship. However, contrary to findings in animals (Romiguier et al., 2014), genetic diversity across butterflies could not be explained by longevity or propagule size (Mackintosh et al., 2019). One may hypothesize that life history indicators of the strength of genetic drift may vary depending on the taxon or the evolutionary scale of interest. Within flies, developmental time is likely unable to capture variation in genetic drift intensity, it would be very useful to collect other life history traits for those species, such as age and size at maturity, fecundity and fertility to test whether any of them could to be a good indicator of genetic drift.

The nearly neutral theory of molecular evolution predicts that selection efficacy on codon usage bias depends on a species’ effective population size (Kimura, 1983; Charlesworth, 2009), suggesting that the variation of this selection strength among close relatives may result from variation in *Ne* since their divergence. Galtier et al. (2018) found evidence for selection on codon usage only in large-*Ne* species of animals, but not in small-*Ne* ones. Evidence for selection on codon usage has previously been reported in *D. melanogaster* (Shields et al., 1988), *but it remains unclear whether effective selection on codon usage is acting in other fly species. Finally, interspecific differences in Ne* and selection on codon usage may be weakly and inconsistently correlated in flies, as it was found in mammals (Kessler and Dean, 2014).

### 3.4 About the correlation between *dN/dS* and TE content

While we found substantial evidence for a positive correlation between the *dN/dS* ratio and the TE content, the signal was dependent on the method used. Using PGLS and last branch *dN/dS* to test this correlation led to a weaker signal than an approach also considering the *dN/dS* of internal branches (using coevol), showing the importance of more integrative methods. Moreover, the signal for a positive correlation between *dN/dS* and TE content also varied according to the gene set used. In particular, the strength of this correlation was stronger for GC3-poor genes and lower for GC3-high genes. This suggests that the variation in evolutionary rates among genes is not independent from their GC3. However, we did not detect a direct relationship between the concatenate GC3 and the coefficient of the correlation between *dN/dS* and TE content. The correlation between *dN/dS* and TE content tends to be more often significant using concatenates of genes homogeneous for their GC3 compared to random set of genes.

This can be related to the fact that an heterogeneity in base composition among genes within a concatenate is expected to alter the codon frequency estimate (which won’t be correct at the gene scale), and lessen the accuracy of the estimation of the global *dN/dS* based on phylogenetic models. Moreover, the GC3 of genes is not completely independent of the coefficient of purifying selection. In particular, highly expressed genes are expected to be enriched in optimal codons due to selection on codon usage, and optimal codons are mainly G- or C-ending in *Drosophila* (Duret and Mouchiroud, 1999). Therefore, high-GC3 genes are expected to evolve under stronger selection than poor-GC3 genes. While *Ne*-reduced species may prevent the fixation of non synonymous deleterious mutations in highly expressed genes (enriched in high-GC3 genes), they may show an accumulation of slightly deleterious mutations in less expressed genes (enriched in poor-GC3 genes). Furthermore, highly expressed genes evolve under selection for codon usage bias, at least in *D. melanogaster* (Duret and Mouchiroud, 1999), meaning that *dS* is not neutral in those genes, and therefore we may question the use of their *dN/dS* to provide an accurate estimate of the effective population size.

Comparative studies were previously conducted among closely related species with different *Ne* associated with changes in various life history traits, and they provide contrasting results (reported on table 3). Most studies that did not find any evidence of TE accumulation associated with a reduction in *Ne* did compare asexual lines or species with sexual ones. However, if a transition to asexuality is expected to induce a drastic reduction in *Ne*, it has other evolutionary implications. In particular, Charlesworth and Langley (1986) proposed that TE transposition rates should reduce due to within lineage transmission in non recombining genomes. In a nutshell, the reduction in transposition in asexuals would override the trend for TE accumulation due to increased genetic drift. The contrasting results among studies comparing eusocial species with noneusocial relatives are more complex to interpret. Eusocial species are expected to harbour reduced *Ne*, and very high levels of recombination were found in social insects (Wilfert et al., 2007). Therefore, TE accumulation is expected in eusocial species, as found in Korb et al. (2015); Chak et al. (2021), but not in Ardila-Garcia et al. (2010); Kapheim et al. (2015); Koshikawa et al. (2008). One possible explanation might lie in the differences in the evolutionary scale. Indeed, compared to bees, snapping shrimps have diverged much more recently, which might have blurred the signal due to confounding factors. An example of a possible factor that prevented the observation of TE accumulation in the reduced-*Ne*-species is a change in the epigenetic TE-silencing mecanisms. If the synonymous divergence is too large, saturation could occur, leading to an underestimation of the *dN/dS* ratio and a least accurate proxy of *Ne*. Alternatively, according to the rate of horizontal transfers of TEs occurring in some species, it is possible that the TE content could be better explained by different invasion histories or variation in the defense mechanisms than by the intensity of the genetic drift. Indeed, we acknowledge that the total TE content is a simple measure resulting from complex interactions of various mechanisms.

**Table 3:**
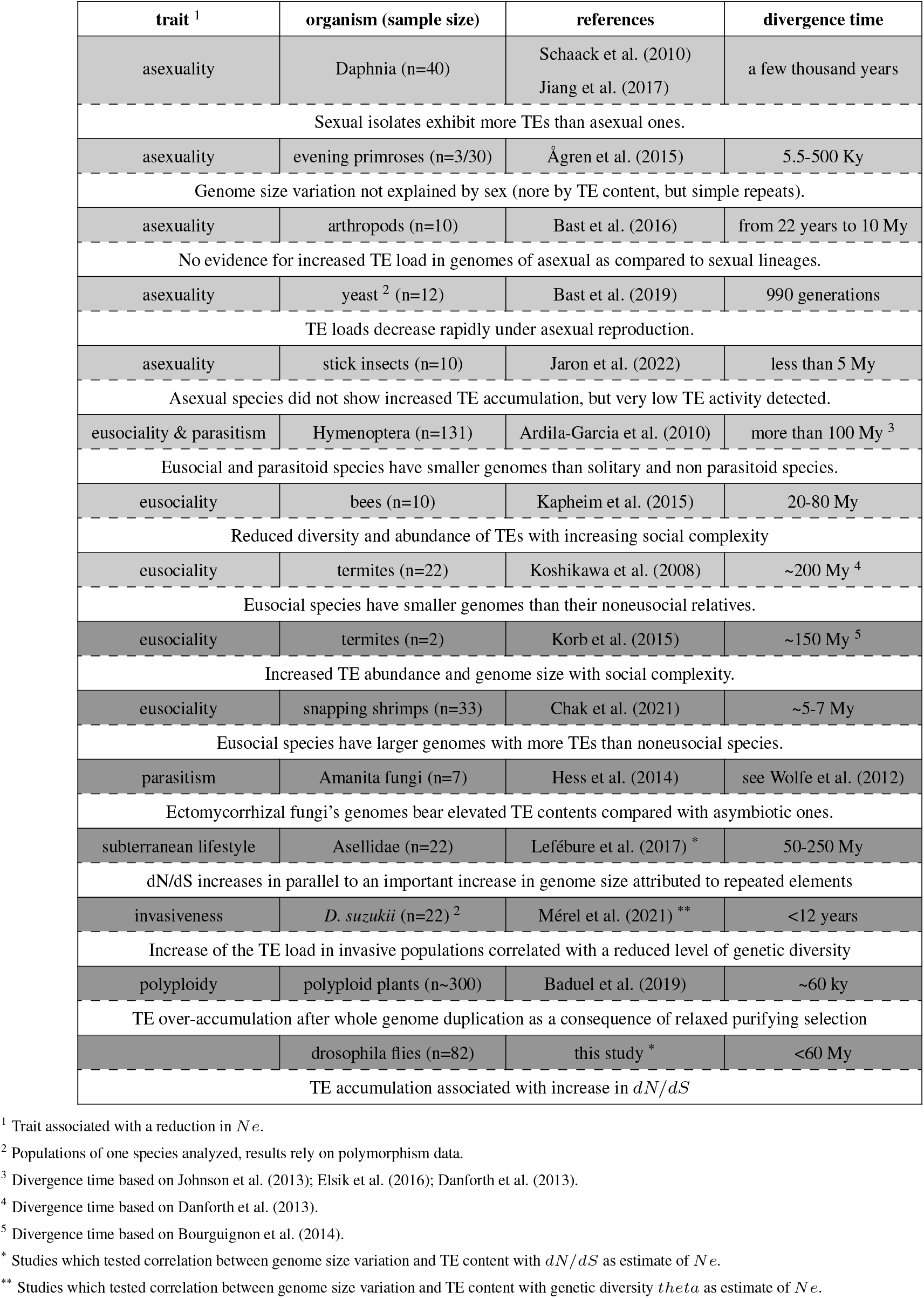
Literature review of genome size and TE content changes resulting from transitions in life history traits associated with a reduction in *Ne*. Studies presented in the lightgray lines did not provide support to the *Mutational Hazard* theory, as they did not find any evidence for accumulation of TEs or increase in genome size in lineages with reduced *Ne*. On the contrary, darker gray lines report studies where accumulation of TEs or increase in genome size has been identified in lineages with reduced *Ne*.

In flies, we found a significant positive correlation between the TE content and the *dN/dS* ratio, that is to say an accumulation of TEs in species with lower *Ne* estimated through their selection efficiency. A similar relationship between TE load and genomic estimates of *Ne* was found in subterranean Asellidae (Lefébure et al., 2017), or at an intraspecific level within *D. suzukii*, where invasive populations showed reduced genetic diversity and increased TE content (Mérel et al., 2021).

### 3.5 Conclusion

Genome size and TE abundance are positively correlated within flies. We found a strong support for a relationship between these two variables and a long term estimate of *Ne* (*dN/dS*), suggesting a substantial effect of relaxed purifying selection on TE proliferation. Support for this correlation was dependant on the gene set used, which is likely to reflect different evolutionary rates between genes. In particular, our results suggest that GC heterogeneity should be controlled for to improve estimates of *dN/dS*. As our knowledge of flies progressively extends beyond *D. melanogaster*, we should soon be able to integrate new variables, such as local recombination rate or gene expression level, in interspecific comparative studies, allowing a better understanding of the complex relationship between relaxed purifying selection and molecular traits.

## 4 Methods

### 4.1 Fly sampling

Genome size, TE content and selection efficacy were studied for 77 drosophilid species (73 *Drosophila*, three *Zaprionus*, and one *Scaptodrosophila* - supplementary table S1). DNA sequencing data used to infer TE content and selection efficacy were retrieved from public databases for 43 species (supplementary table S2,3,5). Flow cytometry estimates were retrieved from public databases for 59 species (supplementary table S4). For cases where sequencing data and/or flow cytometry estimates were not available, flies were collected from stock centers and research labs (supplementary table S7).

The strains were ordered to obtain lines from a single female that was sampled in the wild as recently as possible. Since the study lines were derived from relatively recently sampled females, the genome fixed in these lines should be representative of a genome from a natural population. Moreover isofemale lines, maintained in the lab with small population sizes, should be mostly homozygous, facilitating assembly.

Flies were bred in the laboratory under their optimal living conditions in order to amplify them. As the amplification generations progressed, the adults were frozen after sex determination. In order to sequence only homologous chromosomes, we chose to sequence female individuals (males are heterogametic). For each species, 60 females were frozen to constitute a sufficient stock of biological material. In parallel, for the estimation of genome size by flow cytometry, the heads of 50 females per species were frozen.

### 4.2 Genome size estimations

Genome size estimates were collected from the Animal Genome Size database (Gregory, 2005), retrieved from Hjelmen et al. (2019), or evaluated using flow cytometry (supplementary table S4). One to sixteen heads were crushed in buffer with heads from a control (either *D. melanogaster, D. simulans*, or *D. virilis*). After addition of propidiumiodid results were analyzed using the BD Accuri ^*T M*^ C6 Plus Flow Cytometer. Replicates were performed per fly line, unless biological material was lacking (only one estimate was possible for *D. tristis*, two for *D. algonquin*; otherwise five to ten replicates were done).

### 4.3 DNA extraction and sequencing

The genome of 38 lines was sequenced for assembly (referred as “homemade assemblies”, supplementary table S6). To obtain sufficient quantity of genetic material for sequencing, DNA was extracted from 20 frozen females per species using a Phenol-Chloroform protocol. The DNA treated with RNAse, was checked on gel and then assayed. Sequencing was performed with TruSeq gDNA libraries (inserts of approximately 500 bp) and HiSeq Illumina paired-ends (2×125 pb) (supplementary table S2).

### 4.4 Genome assemblies

For *de novo* genomes, raw reads were filtered and trimmed for quality using UrQt (Modolo and Lerat, 2015). Two alternative strategies of assemblies were tested. The first consists in progressive depth assemblies using IDBA-UD (Peng et al., 2012), to reduce errors in high-depth regions. The second strategy consists in several steps: i) contig assembly using Ray-2.3.1 (Boisvert et al., 2010), with k-mer size varying from 25 to 45, ii) quality estimation using N50, longest contig size, number of contigs >100kb of the 21 obtained contig assemblies, iii) scaffolding of the three best contig assemblies using SOAP *de novo* (Li et al., 2010), with k-mer size ranging from 59 to 73 (with an increment of 2), iv) selection of the best assembly based on quality estimation using N50, longest scaffold size, number of scaffolds >100kb and coverage portion of the genome size. For each of the 38 “homemade” genomes, we therefore obtained two different assemblies (one with the first strategy, one with the second one). Genomes’ completeness was estimated using Diptera single-copy orthologous gene sets from BUSCO v4 (Seppey et al., 2019): we kept the genome assembly maximizing the number of single-copy core genes found (30 were IDBA-UD and eight RAY+SOAP strategy - supplementary table S2). Publicly available genome assemblies analyzed are listed in the supplementary table S3, along with the source, taxon and assembly ID.

### 4.5 Coding sequences identification & Phylogeny

On both “homemade” and public genome assemblies, repeat sequences were identified and masked from the assemblies using RepeatMasker (RepBase database) (Smit et al., 2013). Genes were predicted using AUGUSTUS v3.1, pre-trained on the *D. melanogaster* genome (Stanke and Waack, 2003). The number of protein-coding genes predicted on each assembly (both “homemade” and public) is provided in supplementary tables S2-3).

Prior to gene clustering, we removed isoform variants from the *D. melanogaster* annotation by conserving only the longest one. We clustered amino acid sequences using MMseqs2 (Steinegger and Söding, 2017), with default parameters. For each cluster, we built a profile HMM using the program hmmbuild from the HMMER package (Eddy, 2011). Then, we compared all HMM profiles for homology using the program hmmsearch from the HMMER package. We merged homologous families with a minimum of 30% overlap using SiliX (Miele et al., 2011). We removed from each family sequences that were smaller than 50% of the median length of the families.

For the phylogenetic reconstruction, we conserved only single-copy orthologs families composed of at least 50% of the 82 lineages, resulting in a dataset of 2,242 gene families. Every nucleotide gene family was aligned with MACSE v2.03 (Ranwez et al., 2011)(options -local_realign_init 1 -local_realign_dec 1 -max_refine_iter 25), then sites that were shared by less than 50% of the sequences were removed with Gblocks 0.91b (Castresana, 2000) (options -b1=[Number of sequences] -b2=[Number of sequences] -b3=10 -b4=2 -b5=h). We inferred a maximum likelihood phylogenetic tree from the concatenation of all amino-acids alignments using IQ-Tree version 1.6.9 (Nguyen et al., 2015), for each partition the most likely model was estimated. We computed 1,000 ultrafast bootstraps (UFBoot) and 1,000 SH-like approximate likelihood ratio test (options -bb 1000 -alrt 1000 -bnni). We also reconstructed the species tree using a supertree approach. For this, all the 2,242 gene trees were individually reconstructed using IQ-Tree, the most likely model being estimated for each gene. We then used ASTRAL-III (Zhang et al., 2018) to infer the species supertree from the individual gene trees, while accounting for gene tree discordance.

### 4.6 Estimations of TE content

TE content was evaluated from private and public paired-end reads dataset using dnaPipeTE (Goubert et al., 2015). dnaPipeTE reconstructs consensus of repeated sequences by assembling a subsample of reads of coverage inferior to 1*X*. A read processing was performed prior to this analysis. Low quality positions were removed and size was unified across samples to 100bp using fastx-toolkit (fastq_quality_filter -q 20 -p 80; fastx_trimmer -l 99; v0.0.13). Note that for *Drosophila mauritiana* reads size was only 75bp. Putative bacterial reads were filtered using kraken2 with NCBI bacterial genomes (v2.1.2: Wood et al. (2019)). dnaPipeTE was used on processed reads, as follow: -genome_coverage 0.15, -sample_number 2 (v1.3.1, (Goubert et al., 2015)).

### 4.7 Phylogenetic inertia, correlations between genome size and TE abundances

Phylogenetic inertia was evaluated using Pagel’s λ (phytools v1.5, (Revell, 2012)). Prior to calculations the tree was converted to ultrametric using ete3 (Huerta-Cepas et al., 6 06), species replicates removed, and values were log-transformed except for developmental time. To evaluate correlations between log transformed genome size and TE abundances while controlling for phylogenetic signal we used the software ancov (Lartillot, 2014). ancov, based on a Kalman filtering algorithm, estimates correlation between quantitative traits using a combination of Markov chain Monte Carlo and Bayesian methods. Additionally, we performed Phylogenetic Generalised Least Squares (PGLS) analysis using the caper package (comparative.data(vcv=TRUE), pgls(lambda = 1.0, kappa = 1.0, delta = 1.0), (Orme et al., 2013)).

### 4.8 Developmental time and Strength of Selection

Developmental time from egg to adult at 18°C were retrieved from (Markow and O’Grady, 2005) for 50 species (supplementary table S8). Molecular traits were estimated from concatenates of 50 genes built from the 2,242 single copy orthologs. Prior to concatenation, nucleotide alignments were filtered using HMMCleaner (Di Franco et al., 2019) and Gblocks, using the same parameters as previously but keeping only sites shared by 90% of the sequences. In order to account for intragenomic differences, we first used three concatenates: one including random genes, one including the 50 genes of lowest GC3, and one with the 50 genes of highest GC3. The analysis was then repeated using ten concatenates of random genes, and 16 concatenates of similar GC3. The latters were built to span the overall GC3 distribution without overlap between concatenates (i.e. a gene can not be within two concatenates).

Codon Usage Bias (CUB) metrics were calculated with a R script using the coRdon package (BioinfoHR, 2022). Metrics were first calculated per sequence, and then the median was computed for each concatenate. We chose as CUB metrics the Effective Number of Codons that explicitly accounts for GC bias (ENC’) (Novembre, 2002) and the Synonymous Codon Usage Score (SCUO), an index of deviation from a uniform distribution based on the Shannon entropy (Wan et al., 2004). Both metrics are not strongly correlated with each other (Liu et al., 2018).

For each concatenate, two different estimations of *dN/dS* were performed using coding sequence alignments. First, coevol was used to reconstruct *dN/dS* evolution across the tree (Lartillot and Poujol, 2011) (see also Correlations between traits). Second bio++ libraries were used to estimate a per branch *dN/dS* (Dutheil and Boussau, 2008; Guéguen et al., 2013). Using previously inferred tree topology, the YN98 (F3X4) was used to retrieved the most likely branch lengths, codon frequencies at the root, and substitution model parameters. Then, MapNH was used to estimate *dN* and *dS* (Guéguen and Duret, 2018; *Minin and Suchard*, *2008; Dutheil et al*., *2012)*.

### 4.9 Correlations between traits

Two different methods were used to assess potential correlations between genome size/TE abundances and developmental time, and estimates of the strength of selection. We first used PGLS to test correlations between genome size/repeat content and selection efficacy, using bio++ last-branch *dN/dS*. The other one is implemented in coevol. coevol does not only model *dN/dS* evolution given a tree and alignments, but also the evolution of other traits while assessing correlations between traits (Lartillot and Poujol, 2011). We thus used coevol to evaluate correlation between genome size, TE content, developmental time, codon bias metrics and *dN/dS* (as a proxy of *Ne*). Variables are assuming brownian motion parametrized by a covariance matrix. All parameters are estimated in Bayesian framework using Monte Carlo Markov Chains. The advantage of such an approach is its integrative nature: in a single test, the correlation between all traits is tested across the entire phylogenetic history.

### 4.10 Statistical analysis

All descriptive and inferential statistics were performed using R (R Core Team, 2021). P-values were corrected for multiple testing using Benjamini-Hochberg procedure (Benjamini and Hochberg, 1995).

## 5 Data access

Details about the 82 drosophilid lineages are provideed in supplementary table S1. Public DNA sequencing data accession numbers are specified in supplementary table S2,3,5. Raw sequencing data were deposited in the Sequence Read Archive repository under the BioProject accession number **XXXXXX**. All scripts are available at https://github.com/vpymerel/GND.

## 7 Acknowledgments

This work was performed using the computing facilities of the CC LBBE/PRABI. A.V.H is supported by the Agence Nationale de la Recherche, grant ANR-20-CE02-0008 / NEGA and the CNRS APEGE DynET. We are thankful to M. Bastian and N. Lartillot for their helpful advice using coevol, to Florian Bénitière for computing the link between GC3 and gene expression supporting our postulate that GC3 rich genes are more expressed than GC3 poor ones, and to Laurent Duret and Julien Joseph for their useful comments on the manuscript.

## References

Ågren, J. A., Greiner, S., Johnson, M. T., and Wright, S. I. (2015). No evidence that sex and transposable elements drive genome size variation in evening primroses. Evolution, 69(4):1053–1062.

Ardila-Garcia, A., Umphrey, G., and Gregory, T. (2010). An expansion of the genome size dataset for the insect order hymenoptera, with a first test of parasitism and eusociality as possible constraints. Insect Molecular Biology, 19(3):337–346.

Baduel, P., Quadrana, L., Hunter, B., Bomblies, K., and Colot, V. (2019). Relaxed purifying selection in autopolyploids drives transposable element over-accumulation which provides variants for local adaptation. Nature Communications, 10(1).

Bast, J., Jaron, K. S., Schuseil, D., Roze, D., and Schwander, T. (2019). Asexual reproduction reduces transposable element load in experimental yeast populations. Elife, 8:e48548.

Bast, J., Schaefer, I., Schwander, T., Maraun, M., Scheu, S., and Kraaijeveld, K. (2016). No accumulation of transposable elements in asexual arthropods. Molecular biology and evolution, 33(3):697–706.

Benjamini, Y. and Hochberg, Y. (1995). Controlling the false discovery rate: A practical and powerful approach to multiple testing. Journal of the Royal Statistical Society: Series B (Methodological), 57(1):289–300.

Bioinfo HR (2022). coRdon. original-date: 2018-06-08T08:16:53Z.

Biémont, C. and Vieira, C. (2006). Junk DNA as an evolutionary force. Nature, 443(7111):521–524.

Boisvert, S., Laviolette, F., and Corbeil, J. (2010). Ray: simultaneous assembly of reads from a mix of high-throughput sequencing technologies. Journal of Computational Biology: A Journal of Computational Molecular Cell Biology, 17(11):1519–1533.

Bolívar, P., Guéguen, L., Duret, L., Ellegren, H., and Mugal, C. F. (2019). GC-biased gene conversion conceals the prediction of the nearly neutral theory in avian genomes. Genome Biology, 20(1):5.

Bourguignon, T., Lo, N., Cameron, S. L., Šobotník, J., Hayashi, Y., Shigenobu, S., Watanabe, D., Roisin, Y., Miura, T., and Evans, T. A. (2014). The evolutionary history of termites as inferred from 66 mitochondrial genomes. Molecular biology and evolution, 32(2):406–421.

Castillo, D. M., Mell, J. C., Box, K. S., and Blumenstiel, J. P. (2011). Molecular evolution under increasing transposable element burden in Drosophila: A speed limit on the evolutionary arms race. BMC Evolutionary Biology, 11(1):258.

Castresana, J. (2000). Selection of Conserved Blocks from Multiple Alignments for Their Use in Phylogenetic Analysis. Molecular Biology and Evolution, 17(4):540–552.

Chak, S. T., Harris, S. E., Hultgren, K. M., Jeffery, N. W., and Rubenstein, D. R. (2021). Eusociality in snapping shrimps is associated with larger genomes and an accumulation of transposable elements. Proceedings of the National Academy of Sciences, 118(24):e2025051118.

Charlesworth, B. (2009). Effective population size and patterns of molecular evolution and variation. Nature Reviews Genetics, 10(3):195–205.

Charlesworth, B. and Barton, N. (2004). Genome size: does bigger mean worse? Current Biology, 14(6):R233–R235.

Charlesworth, B. and Langley, C. (1986). The evolution of self-regulated transposition of transposable elements. Genetics, 112(2):359–383.

Danforth, B. N., Cardinal, S., Praz, C., Almeida, E. A., and Michez, D. (2013). The impact of molecular data on our understanding of bee phylogeny and evolution. Annual review of Entomology, 58:57–78.

Di Franco, A., Poujol, R., Baurain, D., and Philippe, H. (2019). Evaluating the usefulness of alignment filtering methods to reduce the impact of errors on evolutionary inferences. BMC Evolutionary Biology, 19(1):1–17.

Duret, L. and Mouchiroud, D. (1999). Expression pattern and, surprisingly, gene length shape codon usage in caenorhabditis, drosophila, and arabidopsis. Proceedings of the National Academy of Sciences, 96(8):4482–4487.

Dutheil, J. and Boussau, B. (2008). Non-homogeneous models of sequence evolution in the Bio++ suite of libraries and programs. BMC Evolutionary Biology, 8(1):255.

Dutheil, J. Y., Galtier, N., Romiguier, J., Douzery, E. J. P., Ranwez, V., and Boussau, B. (2012). Efficient selection of branch-specific models of sequence evolution. Molecular Biology and Evolution, 29(7):1861–1874.

Eddy, S. R. (2011). Accelerated Profile HMM Searches. PLOS Computational Biology, 7(10):e1002195. Publisher: Public Library of Science.

Elliott, T. A. and Gregory, T. R. (2015). What’s in a genome? the c-value enigma and the evolution of eukaryotic genome content. Philosophical Transactions of the Royal Society B: Biological Sciences, 370(1678):20140331.

Elsik, C. G., Tayal, A., Diesh, C. M., Unni, D. R., Emery, M. L., Nguyen, H. N., and Hagen, D. E. (2016). Hymenoptera genome database: integrating genome annotations in hymenopteramine. Nucleic acids research, 44(D1):D793–D800.

Felsenstein, J. (1985). Phylogenies and the comparative method. The American Naturalist, 125(1):1–15.

Fernández, P., Amice, R., Bruy, D., Christenhusz, M. J., Leitch, I. J., Leitch, A. L., Pokorny, L., Hidalgo, O., and Pellicer, J. (2024). A 160 gbp fork fern genome shatters size record for eukaryotes. iScience.

Fierst, J. L., Willis, J. H., Thomas, C. G., Wang, W., Reynolds, R. M., Ahearne, T. E., Cutter, A. D., and Phillips, P. C. (2015). Reproductive mode and the evolution of genome size and structure in caenorhabditis nematodes. PLoS genetics, 11(6):e1005323.

Flynn, J. M., Long, M., Wing, R. A., and Clark, A. G. (2020). Evolutionary dynamics of abundant 7-bp satellites in the genome of drosophila virilis. Molecular biology and evolution, 37(5):1362–1375.

Galtier, N., Roux, C., Rousselle, M., Romiguier, J., Figuet, E., Glémin, S., Bierne, N., and Duret, L. (2018). Codon usage bias in animals: disentangling the effects of natural selection, effective population size, and gc-biased gene conversion. Molecular biology and evolution, 35(5):1092–1103.

Goubert, C., Modolo, L., Vieira, C., ValienteMoro, C., Mavingui, P., and Boulesteix, M. (2015). De Novo Assembly and Annotation of the Asian Tiger Mosquito (Aedes albopictus) Repeatome with dnaPipeTE from Raw Genomic Reads and Comparative Analysis with the Yellow Fever Mosquito (Aedes aegypti). Genome Biology and Evolution, 7(4):1192–1205.

Gregory, T. R. (2005). Animal genome size database. http://www.genomesize.com/search.php. Accessed: 2021-06-07.

Guéguen, L. and Duret, L. (2018). Unbiased Estimate of Synonymous and Nonsynonymous Substitution Rates with Nonstationary Base Composition. Molecular Biology and Evolution, 35(3):734–742.

Guéguen, L., Gaillard, S., Boussau, B., Gouy, M., Groussin, M., Rochette, N. C., Bigot, T., Fournier, D., Pouyet, F., Cahais, V., Bernard, A., Scornavacca, C., Nabholz, B., Haudry, A., Dachary, L., Galtier, N., Belkhir, K., and Dutheil, J. Y. (2013). Bio++: efficient extensible libraries and tools for computational molecular evolution. Molecular Biology and Evolution, 30(8):1745–1750.

Haddrill, P. R., Loewe, L., and Charlesworth, B. (2010). Estimating the parameters of selection on nonsynonymous mutations in drosophila pseudoobscura and d. miranda. Genetics, 185(4):1381–1396.

Hess, J., Skrede, I., Wolfe, B. E., LaButti, K., Ohm, R. A., Grigoriev, I. V., and Pringle, A. (2014). Transposable element dynamics among asymbiotic and ectomycorrhizal amanita fungi. Genome Biology and Evolution, 6(7):1564–1578.

Hjelmen, C. E., Blackmon, H., Holmes, V. R., Burrus, C. G., and Johnston, J. S. (2019). Genome Size Evolution Differs Between Drosophila Subgenera with Striking Differences in Male and Female Genome Size in Sophophora. G3: Genes, Genomes, Genetics, 9(10):3167–3179. Publisher: G3: Genes, Genomes, Genetics Section: Investigations.

Huerta-Cepas, J., Serra, F., and Bork, P. (2016-06). ETE 3: Reconstruction, analysis, and visualization of phylogenomic data. 33(6):1635–1638.

Jaron, K. S., Parker, D. J., Anselmetti, Y., Tran Van, P., Bast, J., Dumas, Z., Figuet, E., François, C. M., Hayward, K., Rossier, V., et al. (2022). Convergent consequences of parthenogenesis on stick insect genomes. Science advances, 8(8):eabg3842.

Jiang, X., Tang, H., Ye, Z., and Lynch, M. (2017). Insertion polymorphisms of mobile genetic elements in sexual and asexual populations of daphnia pulex. Genome biology and evolution, 9(2):362–374.

Johnson, B. R., Borowiec, M. L., Chiu, J. C., Lee, E. K., Atallah, J., and Ward, P. S. (2013). Phylogenomics resolves evolutionary relationships among ants, bees, and wasps. Current Biology, 23(20):2058–2062.

Kapheim, K. M., Pan, H., Li, C., Salzberg, S. L., Puiu, D., Magoc, T., Robertson, H. M., Hudson, M. E., Venkat, A., Fischman, B. J., et al. (2015). Genomic signatures of evolutionary transitions from solitary to group living. Science, 348(6239):1139–1143.

Kapitonov, V. V. and Jurka, J. (2008). A universal classification of eukaryotic transposable elements implemented in Repbase. Nature Reviews. Genetics, 9(5):411–412; author reply 414.

Kessler, M. D. and Dean, M. D. (2014). Effective population size does not predict codon usage bias in mammals. Ecology and evolution, 4(20):3887–3900.

Kidwell, M. G. (2002). Transposable elements and the evolution of genome size in eukaryotes. Genetica, 115:49–63.

Kim, B. Y., Wang, J. R., Miller, D. E., Barmina, O., Delaney, E., Thompson, A., Comeault, A. A., Peede, D., D’Agostino, E. R., Pelaez, J., Aguilar, J. M., Haji, D., Matsunaga, T., Armstrong, E. E., Zych, M., Ogawa, Y., Stamenković-Radak, M., Jelić, M., Veselinović, M. S., Tanasković, M., Erić, P., Gao, J.-J., Katoh, T. K., Toda, M. J., Watabe, H., Watada, M., Davis, J. S., Moyle, L. C., Manoli, G., Bertolini, E., Košt’ál, V., Hawley, R. S., Takahashi, A., Jones, C. D., Price, D. K., Whiteman, N., Kopp, A., Matute, D. R., and Petrov, D. A. (2021). Highly contiguous assemblies of 101 drosophilid genomes. eLife, 10:e66405. Publisher: eLife Sciences Publications, Ltd.

Kimura, M. (1983). The neutral theory of molecular evolution. Cambridge University Press.

Korb, J., Poulsen, M., Hu, H., Li, C., Boomsma, J. J., Zhang, G., and Liebig, J. (2015). A genomic comparison of two termites with different social complexity. Frontiers in Genetics, 6:9.

Koshikawa, S., Miyazaki, S., Cornette, R., Matsumoto, T., and Miura, T. (2008). Genome size of termites (insecta, dictyoptera, isoptera) and wood roaches (insecta, dictyoptera, cryptocercidae). Naturwissenschaften, 95:859–867.

Lartillot, N. (2014). A phylogenetic Kalman filter for ancestral trait reconstruction using molecular data. Bioinformatics, 30(4):488–496.

Lartillot, N. and Delsuc, F. (2012). Joint reconstruction of divergence times and life-history evolution in placental mammals using a phylogenetic covariance model. Evolution, 66(6):1773–1787.

Lartillot, N. and Poujol, R. (2011). A phylogenetic model for investigating correlated evolution of substitution rates and continuous phenotypic characters. Molecular Biology and Evolution, 28(1):729–744.

Lefébure, T., Morvan, C., Malard, F., François, C., Konecny-Dupré, L., Guéguen, L., Weiss-Gayet, M., Seguin-Orlando, A., Ermini, L., Sarkissian, C. D., Charrier, N. P., Eme, D., Mermillod-Blondin, F., Duret, L., Vieira, C., Orlando, L., and Douady, C. J. (2017). Less effective selection leads to larger genomes. Genome Research, 27(6):1016–1028.

Li, R., Zhu, H., Ruan, J., Qian, W., Fang, X., Shi, Z., Li, Y., Li, S., Shan, G., Kristiansen, K., Li, S., Yang, H., Wang, J., and Wang, J. (2010). De novo assembly of human genomes with massively parallel short read sequencing. Genome Research, 20(2):265–272.

Liu, S. S., Hockenberry, A. J., Jewett, M. C., and Amaral, L. A. N. (2018). A novel framework for evaluating the performance of codon usage bias metrics. Journal of the Royal Society Interface, 15(138):20170667.

Lynch, M. and Conery, J. S. (2003). The Origins of Genome Complexity. Science, 302(5649):1401–1404.

Mackintosh, A., Laetsch, D. R., Hayward, A., Charlesworth, B., Waterfall, M., Vila, R., and Lohse, K. (2019). The determinants of genetic diversity in butterflies. Nature communications, 10(1):3466.

Markow, T. A. and O’Grady, P. (2005). Chapter: Life history variation. In Drosophila: a guide to species identification and use. Elsevier.

Martins, E. P. and Hansen, T. F. (1997). Phylogenies and the comparative method: a general approach to incorporating phylogenetic information into the analysis of interspecific data. The American Naturalist, 149(4):646–667.

McVean, G. A. and Charlesworth, B. (1999). A population genetic model for the evolution of synonymous codon usage: patterns and predictions. Genetics Research, 74(2):145–158.

Mérel, V., Boulesteix, M., Fablet, M., and Vieira, C. (2020). Transposable elements in drosophila. Mobile DNA, 11:1–20.

Mérel, V., Gibert, P., Buch, I., Rodriguez Rada, V., Estoup, A., Gautier, M., Fablet, M., Boulesteix, M., and Vieira, C. (2021). The worldwide invasion of drosophila suzukii is accompanied by a large increase of transposable element load and a small number of putatively adaptive insertions. Molecular biology and evolution, 38(10):4252–4267.

Miele, V., Penel, S., and Duret, L. (2011). Ultra-fast sequence clustering from similarity networks with SiLiX. BMC Bioinformatics, 12(1):116.

Minin, V. N. and Suchard, M. A. (2008). Fast, accurate and simulation-free stochastic mapping. Philosophical Transactions of the Royal Society B: Biological Sciences, 363(1512):3985–3995.

Modolo, L. and Lerat, E. (2015). UrQt: an efficient software for the Unsupervised Quality trimming of NGS data. BMC Bioinformatics, 16(1):137.

Nguyen, L.-T., Schmidt, H. A., von Haeseler, A., and Minh, B. Q. (2015). IQ-TREE: a fast and effective stochastic algorithm for estimating maximum-likelihood phylogenies. Molecular Biology and Evolution, 32(1):268–274.

Novembre, J. A. (2002). Accounting for background nucleotide composition when measuring codon usage bias. Molecular Biology and Evolution, 19(8):1390–1394.

Obbard, D. J., Maclennan, J., Kim, K.-W., Rambaut, A., O’Grady, P. M., and Jiggins, F. M. (2012). Estimating Divergence Dates and Substitution Rates in the Drosophila Phylogeny. Molecular Biology and Evolution, 29(11):3459–3473.

Orgel, L. E. and Crick, F. H. C. (1980). Selfish DNA: the ultimate parasite. Nature, 284(5757):604–607.

Orme, D., Freckleton, R., Thomas, G., Petzoldt, T., Fritz, S., Isaac, N., and Pearse, W. (2013). The caper package: comparative analysis of phylogenetics and evolution in r. R package version, 5(2):1–36.

Pagel, M. D. and Harvey, P. H. (1988). Recent developments in the analysis of comparative data. The Quarterly Review of Biology, 63(4):413–440.

Parvathy, S. T., Udayasuriyan, V., and Bhadana, V. (2022). Codon usage bias. Molecular biology reports, 49(1):539–565.

Peng, Y., Leung, H. C. M., Yiu, S. M., and Chin, F. Y. L. (2012). IDBA-UD: a de novo assembler for single-cell and metagenomic sequencing data with highly uneven depth. Bioinformatics, 28(11):1420–1428.

Pollak, E. (1987). On the theory of partially inbreeding finite populations. i. partial selfing. Genetics, 117(2):353–360.

R Core Team (2021). R: A Language and Environment for Statistical Computing. R Foundation for Statistical Computing, Vienna, Austria.

Ranwez, V., Harispe, S., Delsuc, F., and Douzery, E. J. P. (2011). MACSE: Multiple Alignment of Coding SEquences Accounting for Frameshifts and Stop Codons. PLOS ONE, 6(9):e22594. Publisher: Public Library of Science.

Revell, L. J. (2012). phytools: An R package for phylogenetic comparative biology (and other things). Methods in Ecology and Evolution, 3:217–223.

Robinson, D. F. and Foulds, L. R. (1981). Comparison of phylogenetic trees. Mathematical Biosciences, 53(1):131–147.

Romiguier, J., Lourenco, J., Gayral, P., Faivre, N., Weinert, L. A., Ravel, S., Ballenghien, M., Cahais, V., Bernard, A., Loire, E., et al. (2014). Population genomics of eusocial insects: the costs of a vertebrate-like effective population size. Journal of Evolutionary Biology, 27(3):593–603.

Schaack, S., Pritham, E. J., Wolf, A., and Lynch, M. (2010). Dna transposon dynamics in populations of daphnia pulex with and without sex. Proceedings of the Royal Society B: Biological Sciences, 277(1692):2381–2387.

Schoch, C. L., Ciufo, S., Domrachev, M., Hotton, C. L., Kannan, S., Khovanskaya, R., Leipe, D., Mcveigh, R., O’Neill, K., Robbertse, B., Sharma, S., Soussov, V., Sullivan, J. P., Sun, L., Turner, S., and Karsch-Mizrachi, I. (2020). NCBI Taxonomy: a comprehensive update on curation, resources and tools. Database: The Journal of Biological Databases and Curation, 2020:baaa062.

Seppey, M., Manni, M., and Zdobnov, E. M. (2019). BUSCO: Assessing Genome Assembly and Annotation Completeness. Methods in Molecular Biology (Clifton, N.J.), 1962:227–245.

Sessegolo, C., Burlet, N., and Haudry, A. (2016). Strong phylogenetic inertia on genome size and transposable element content among 26 species of flies. Biology Letters, 12(8):20160407.

Shields, D. C., Sharp, P. M., Higgins, D. G., and Wright, F. (1988). “ silent” sites in drosophila genes are not neutral: evidence of selection among synonymous codons. Molecular biology and evolution, 5(6):704–716.

Smit, A., Hubley, R., and Green, P. (2013). RepeatMasker Home Page.

Stanke, M. and Waack, S. (2003). Gene prediction with a hidden Markov model and a new intron submodel. Bioinformatics, 19(suppl_2):ii215–ii225.

Steinegger, M. and Söding, J. (2017). MMseqs2 enables sensitive protein sequence searching for the analysis of massive data sets. Nature Biotechnology, 35(11):1026–1028. Number: 11 Publisher: Nature Publishing Group.

Sun, C., Shepard, D. B., Chong, R. A., López Arriaza, J., Hall, K., Castoe, T. A., Feschotte, C., Pollock, D. D., and Mueller, R. L. (2012). LTR Retrotransposons Contribute to Genomic Gigantism in Plethodontid Salamanders. Genome Biology and Evolution, 4(2):168–183.

Suvorov, A., Kim, B. Y., Wang, J., Armstrong, E. E., Peede, D., D’Agostino, E. R. R., Price, D. K., Wadell, P., Lang, M., Courtier-Orgogozo, V., David, J. R., Petrov, D., Matute, D. R., Schrider, D. R., and Comeault, A. A. (2020). Widespread introgression across a phylogeny of 155 Drosophila genomes. bioRxiv, page 2020.12.14.422758. Publisher: Cold Spring Harbor Laboratory Section: New Results.

Tenaillon, M. I., Hollister, J. D., and Gaut, B. S. (2010). A triptych of the evolution of plant transposable elements. Trends in Plant Science, 15(8):471–478.

Van Der Linde, K., Houle, D., Spicer, G. S., and Steppan, S. J. (2010). A supermatrix-based molecular phylogeny of the family drosophilidae. Genetics research, 92(1):25–38.

Wan, X.-F., Xu, D., Kleinhofs, A., and Zhou, J. (2004). Quantitative relationship between synonymous codon usage bias and GC composition across unicellular genomes. BMC Evolutionary Biology, 4:19.

Whitney, K. D., Garland, T., and Jr (2010). Did Genetic Drift Drive Increases in Genome Complexity? PLoS Genetics, 6(8).

Wilfert, L., Gadau, J., and Schmid-Hempel, P. (2007). Variation in genomic recombination rates among animal taxa and the case of social insects. Heredity, 98(4):189–197.

Wolfe, B. E., Tulloss, R. E., and Pringle, A. (2012). The irreversible loss of a decomposition pathway marks the single origin of an ectomycorrhizal symbiosis. PLOS ONE, 7(7):e39597.

Wood, D. E., Lu, J., and Langmead, B. (2019). Improved metagenomic analysis with Kraken 2. Genome Biology, 20(1):257.

Zhang, C., Rabiee, M., Sayyari, E., and Mirarab, S. (2018). ASTRAL-III: polynomial time species tree reconstruction from partially resolved gene trees. BMC Bioinformatics, 19(6):153.

